# Susceptibility trends of zoliflodacin against multidrug-resistant *Neisseria gonorrhoeae* clinical isolates in Nanjing, China (2014-2018)

**DOI:** 10.1101/2020.05.04.078055

**Authors:** Wenjing Le, Xiaohong Su, Xiangdi Lou, Xuechun Li, Xiangdong Gong, Baoxi Wang, Caroline A. Genco, John P. Mueller, Peter A. Rice

## Abstract

Previously, we reported potent activity of a novel spiropyrimidinetrione, zoliflodacin, against *N. gonorrhoeae* isolates from symptomatic men in Nanjing, China, collected in 2013. Here, we investigated trends of susceptibilities of zoliflodacin in 986 gonococcal isolates collected from men between 2014 and 2018. *N. gonorrhoeae* isolates were tested for susceptibility to zoliflodacin and seven other antibiotics. Mutations in *gyrA, gyrB, parC* and *parE* genes were determined by PCR and DNA sequencing. The MIC of zoliflodacin for *N. gonorrhoeae* ranged from ≤0.002 to 0.25 mg/L; the overall MIC_50_s and MIC_90_s were 0.06 mg/L and 0.125mg/L in 2018, increasing two-fold from 2014. However, the percent of isolates with lower zoliflodacin MICs declined in each year sequentially while the percent with higher MICs increased yearly (P≤0.00001). All isolates were susceptible to spectinomycin but resistant to ciprofloxacin (MIC ≥1 μg/ml); 21.2% (209/986) were resistant to azithromycin (≥1 μg/ml), 43.4% (428/986) were penicillinase-producing (PPNG), 26.9% (265/986) tetracycline-resistant (TRNG) and 19.4% (191/986) were multi-drug resistant (MDR) isolates. Among 143 isolates with higher zoliflodacin MICs (0.125-0.25 mg/L), all had quinolone resistance associated double or triple mutations in *gyrA*; 139/143 (97.2%) also had mutations in *parC*. There were no D429N/A and/or K450T mutations in GyrB identified in the 143 isolates with higher zoliflodacin MICs; a S467N mutation in GyrB was identified in one isolate. We report that zoliflodacin has excellent *in vitro* activity against clinical gonococcal isolates, including those with high-level resistance to ciprofloxacin, azithromycin and extended spectrum cephalosporins.

## INTRODUCTION

*Neisseria gonorrhoeae*, the causative agent of the sexually transmitted infection gonorrhea, has developed resistance to all previously recommended antimicrobial agents for treatment, including sulfonamides, penicillins, tetracyclines and fluoroquinolones^[1]^. Currently, dual antimicrobial therapy with ceftriaxone 250 mg or cefixime 400 mg plus azithromycin 1g is recommended as first-line treatment of uncomplicated gonorrhea by the World Health Organization (WHO)^[2]^ and ceftriaxone plus azithromycin by the U. S. Centers for Disease Control and Prevention (CDC)^[3]^. Resistance to extended-spectrum cephalosporin (ESCs) and azithromycin is increasing worldwide. Gonococcal isolates with decreased susceptibility to cefixime and/or ceftriaxone have been reported in China^[4]^, Japan^[5]^, Australia^[6]^, European countries^[7]^ and the United States^[8]^ and isolates with high-level resistance to ceftriaxone have been identified in Japan, Australia, France, Spain, Denmark, Canada Ireland and China ^[9,10,11]^. The reported prevalence of azithromycin-resistant *N. gonorrhoeae* isolates is 18.6% in China^[4]^, 14.5% in Japan^[5]^, 6.2% in Australia ^[6]^, 7.5% in 25 European countries ^[7]^, 4.6% in the United States ^[8]^, and 6.1% in Western Africa^[12]^. The first documented case that failed treatment with the recommended dual therapy was reported from the UK in 2016 ^[13]^ and the first gonococcal isolates (the A2543 clone) with combined ceftriaxone plus high-level azithromycin resistance were identified in the UK^[14]^ and Australia^[15]^ in 2018.

Increased antimicrobial resistance (AMR) in *N. gonorrhoeae* poses an emerging global public health threat of untreatable gonococcal infections.New oral antimicrobial agents with activity against *N. gonorrhoeae* are needed urgently. WHO includes *N. gonorrhoeae* on its list of “priority pathogens” that require new antibiotics for treatment^[16]^ and the U.S. CDC has designated drug-resistant *N. gonorrhoeae* as an urgent threat^[17]^. Zoliflodacin (also known as AZD0914 and ETX0914) is a novel spiropyrimidinetrione bacterial DNA gyrase /topoisomerase inhibitor with broad-spectrum *in vitro* activity against gram-positive and fastidious gram-negative organisms, including *N. gonorrhoeae*.^[18,19]^. A recent multicenter, randomized, phase 2 clinical trial^[20]^ confirmed the efficacy and safety of a single dose of 2 g or 3 g of oral zoliflodacin compared with 500 mg of intramuscular ceftriaxone for the treatment of uncomplicated gonococcal infections. We showed previously that zoliflodacin was highly effective against clinical isolates of *N. gonorrhoeae in vitro*, including high-level ciprofloxacin-resistant and multidrug resistant isolates, collected in 2013 in Nanjing, China^[21]^. Here, *in vitro* activities and trends of zoliflodacin susceptibilities were determined for clinical gonococcal isolates (including multidrug resistant isolates), collected between 2014 and 2018 in Nanjing. Mutations in the quinolone-resistance-determinant regions (QRDRs) of *gyrA, parC, gyrB* and *parE*genes in isolates with higher MICs (0.125mg/L and 0.25mg/L) to zoliflodacin were also determined.

## RESULTS

### Susceptibilities to zoliflodacin and other antimicrobials

Susceptibilities (MICs) of *N. gonorrhoeae* to zoliflodacin and seven antimicrobials previously or currently used for the treatment of gonorrhea are summarized for the 986 clinical isolates in Table 1. All isolates except one were inhibited by ≤0.125 mg/L of zoliflodacin (the remaining isolate had an MIC of 0.25mg/L). MICs to zoliflodacin ranged from ≤0.002 to 0.25mg/L overall, with an MIC_50_ and MIC_90_ of 0.06 mg/L and 0.125 mg/L, respectively. 143 (14.5%) isolates had zoliflodacin MICs of 0.125-0.25 mg/L. The percent of isolates with an MIC of 0.03 mg/L to zolifodacin declined in each year sequentially (χ^2^= 82.237, P=0.000) while the percent with MICs of 0.06 and 0.125 mg/L increased correspondingly (χ^2^= 20.739 and 41.717, respectively; P≤ 0.00001; Chi square test for linear trend), shown in Figure 1. Overall, the proportion of isolates with zoliflodacin MIC 0.125-0.25 mg/L increased from 3.1% (6/197) in 2014 to 23.0% (47/204) in 2018 (χ^2^= 43.112, P<0.0001).

**Table 1.** Susceptibilities and MICs of zoliflodacin and seven antimicrobials previously or currently used for treatment of gonorrhea against 986 clinical *N. gonorrhoeae* isolates.

| Antimicrobial | No. (%) |  |  | MIC (mg/L) |  |  |
| --- | --- | --- | --- | --- | --- | --- |
|  | Susceptible | Intermediate | Resistant | Range | MIC<sub>50</sub> | MIC<sub>90</sub> |
| zoliflodacin | 986 (100) |  |  | ≤0.002 to 0.25 | 0.06 | 0.125 |
| Penicillin G | 0 | 171 (17.3) | 815 (82.7) | 0.125 to ≥16 | 4 | ≥16 |
| tetracycline | 4 (0.4) | 150 (15.2) | 832 (84.4) | ≤0.125 to ≥32 | 2 | ≥32 |
| ciprofloxacin | 0 | 0 | 986 (100) | 1 to ≥ 16 | ≥16 | ≥16 |
| azithromycin | 551(55.9) | 226 (22.9) | 209 (21.2) | ≤0.015 to ≥2048 | 0.5 | 4 |
| spectinomycin | 986(100) | 0 | 0 | ≤4 to 32 | 32 | 32 |
| cefixime | 948(96.1) | - | 38 (3.9) | ≤0.002 to >2 | 0.03 | 0.25 |
| ceftriaxone | 984(99.8) | - | 2 (0.2) | ≤0.002 to 1 | 0.03 | 0.125 |

MIC: minimum inhibitory concentration

**Figure 1.**
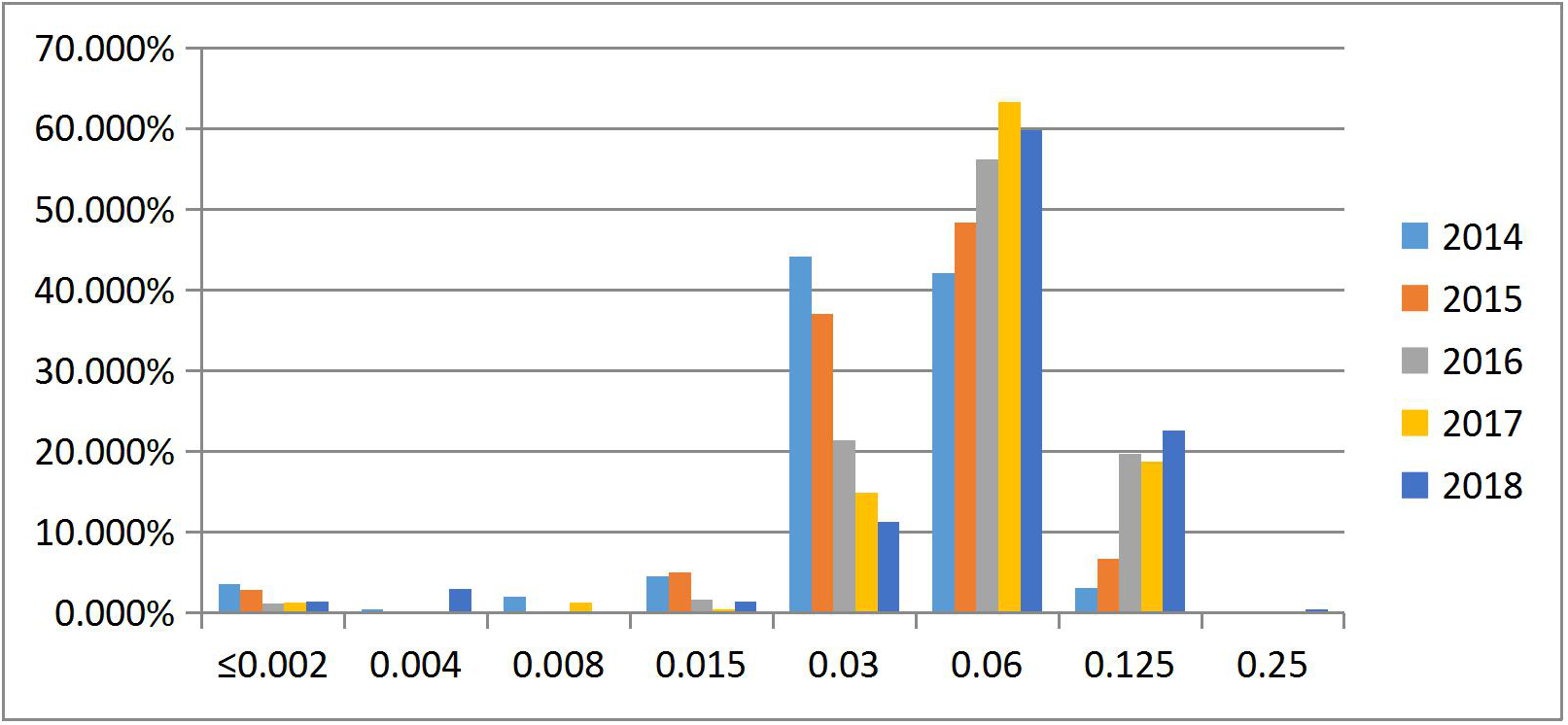
MIC distributions of zoliflodacin for 986 clinical *N. gonorrhoeae* isolates (2014-2018).

All 986 isolates were resistant to ciprofloxacin; 777 (78.8%) showed high level resistance (≥16 mg/L)^[22]^. During the five year study period, the annual percentage of ciprofloxacin resistant isolates at each MIC point (from 1 mg/L to ≥16mg/L) did not shift in either direction in the 5-year period. MICs of gonococcal isolates for zoliflodacin were lower than ciprofloxacin (P<0.0001), with a median difference of atleast 267-fold. Four hundred and twenty eight isolates (43.4%) were PPNG and 265 (26.9%) were TRNG. The percent of penicillin-resistant isolates increased from 70% to 86.3% over the five years (χ^2^= 17.641, P< 0.0001). Although all isolates were susceptible to spectinomycin, the percent of isolates with lower spectinomycin MICs (8 mg/L and 16 mg/L) declined (χ^2^= 16.35 and 93.71, P=0.0001 and P< 0.0001, respectively) while the percent with higher MICs (32mg/L) increased over the five years (χ^2^= 112.514, P<0.0001).

Two hundred and nine (21.2%) isolates were resistant to azithromycin (MIC ≥ 1mg/L), and 62 (6.3%) displayed high-level resistance (MIC ≥256 mg/L). The percent of isolates with lower azithromycin MICs (0.06 mg/L and 0.125mg/L) increased over the five year(χ^2^= 16.916 and 22.099, respectively; P< 0.0001) while the percent with higher MICs (0.5mg/L and ≥1024 mg/L) declined yearly (χ^2^= 15.403 and 12.268, respectively; P<0.001). Overall, the percent of azithromycin-resistant isolates (MIC ≥ 1mg/L) decreased from 27.9% to 15.2% over the five years and the percent of azithromycin-susceptible isolates increased from 72.1% to 84.8% (χ2 = 14.618, P< 0.001). One hundred and fifty eight isolates (15.2%) exhibited decreased susceptibility (MIC 0.125-0.25 mg/L, n=156) or resistance (MIC = 1mg/L, n=2) to ceftriaxone, and 102 isolates (10.1%) displayed decreased susceptibility (MIC 0.25mg/L, n=64) or resistance (MIC 0.5mg/ L, n=36; MIC≥2mg/L, n=2) to cefixime. The percent of isolates with lower ceftriaxone MICs (0.03mg/L) declined in each year sequentially (χ^2^= 10.512, P< 0.01) while the percent with higher MICs (0.06mg/L and 0.125 mg/L) increased yearly (χ^2^= 10.18 and 4.231, P<0.01 and P<0.05, respectively).

The percent of isolates with lower cefixime MICs (0.015 mg/L and 0.03 mg/L) declined (χ^2^= 23.324 and 10.734, P<0.001 and P<0.01, respectively) while the percent with higher MICs (0.06-0.5mg/L) increased over the five years (χ^2^= 10.734, 8.68, 14.683 and 20.056, P<0.05, ~P<0.0001, respectively). One hundred ninety one (19.4%) isolates showed multidrug resistance (MDR). The proportion of MDR isolates increased from 7.1% in 2014 to 27% in 2016, then decreased to 21.1% in 2018 (χ^2^= 12.82, P=0.00034). The two MDR isolates with high level resistance to ceftriaxone (MIC 1.0 mg/L), cefixime (MIC ≥ 2.0 mg / L), ciprofloxacin (MIC ≥ 16mg/L), penicillin (MIC 4 mg/L) and tetracycline (MIC 4mg/L) had low zoliflodacin MIC values (0.03 and 0.06 mg/L, respectively).

### Characterization of amino acid substitutions in GyrA, GyrB, ParC and ParE

One hundred forty three gonococcal isolates with higher MICs (0.125-0.25 mg/L) to zoliflodacin were also ciprofloxacin-resistant (MICs 4 to ≥ 16 mg/L). All isolates had double or triple mutations in the *gyrA* gene. Both S91F and D95A/G/N/Y amino acid substitution in GyrA were identified in the 143 isolates and 16 (11.2%) of these isolates also had an additional A92P amino acid substitution in GyrA. ParC substitutions were observed in 97.2% of the isolates. Single, double and triple ParC substitutions were identified in 114 (79.7%), 22 (15.4%) and 3 (2.1%) of the 143 isolates, respectively. Ninety (62.9%) had a single substitution with either S87R, S87N, S87I or S87C. Nineteen (13.3%) had a D86N substitution and 5 (3.5%) an E91G substitution. The most common double substitutions in ParC were S87R plus S88P (10.7%), followed by G85D plus S87R (4.2%); S87N plus E91G (1.4%); S87R plus G120R (1.4%); D86N plus S87I (0.7%); and G85C plus A89T (0.7%). Three isolates had the same triple substitutions (S87R, A123V and A129V). A89T, G120R, A123V and A129V mutations in ParC are newly described here. GyrB substitutions/insertions were identified in four isolates (two with V470I substitutions, one with a S467N substitution and one with an A insertion at 480[480A1]. All four isolates had an MIC value of 0.125 mg/L for zoliflodacin and 4 mg/L or greater for ciprofloxacin. Amino acid substitutions in ParE were identified in 57 isolates (39.9%). The most common single substitution in ParE was D437N (33 isolates [23.1%]), followed by P456S (22 isolates [15.4%]), D437S (1 isolate [0.7%]) and P469L (1isolate [0.7%]). Among the 57 isolates with mutations in *parE*, 53 (93%) had accompanying double *gyrA* and single or double *parC* mutations; the MICs for ciprofloxacin were 8 mg/L (n=3) and ≥16 mg/L (n=50). Table 2 shows the amino acid substitutions in GyrA, GyrB, ParC and Par E that result from the corresponding gene mutations in 143 gonococcal isolates with zoliflodacin MIC ≥0.125 mg/L.

**Table 2.** Amino acid substitutions in GyrA, GyrB, ParC and ParE in 143 isolates with zoliflodacin MIC 0.125-0.25mg/L

| GyrA |  | GyrB |  | ParC |  | ParE |  |
| --- | --- | --- | --- | --- | --- | --- | --- |
| Mutated sites | No. of isolates | Mutated sites | No. of isolates | Mutated sites | No. of isolates | Mutated sites | No. of isolates |
| S91F | 143 | S467N | 1 | G85C | 1 | D437N | 33 |
| A92P | 16 | V470I | 2 | G85D | 6 | D437H | 1 |
| D95N | 24 | +480A | 1 | D86N | 20 | P456S | 22 |
| D95Y | 18 |  |  | S87R | 68 | P469L | 1 |
| D95G | 35 |  |  | S87C | 5 |  |  |
| D95A | 66 |  |  | S87I | 17 |  |  |
|  |  |  |  | S87N | 24 |  |  |
|  |  |  |  | S88P | 10 |  |  |
|  |  |  |  | A89T | 1 |  |  |
|  |  |  |  | E91G | 7 |  |  |
|  |  |  |  | G120R | 2 |  |  |
|  |  |  |  | A123V | 3 |  |  |
|  |  |  |  | A129V | 3 |  |  |

## DISCUSSION

We determined susceptibility trends in *in vitro* antibacterial activity of zoliflodacin and seven other antimicrobial agents against 986 clinical gonococcal isolates collected over a five-year period (2014-2018). The 986 gonococcal isolates were susceptible to zoliflodacin and all were resistant to ciprofloxacin, Nearly a quarter were resistant to azithromycin or were TRNG isolates. Greater than 40% were PPNG isolates and just under 20% were MDR isolates. All 986 isolates had zoliflodacin MICs below the breakpoint (MIC ≥ 0.5mg/L) that have been proposed, guided by clinical efficacy ^[20]^. Similar to other reports ^[19,23]^, zoliflodacin exhibited an MIC range of 0.002 to 0.25 mg/L and there was no correlation between zoliflodaxin MICs at the upper end of the MIC range and ciprofloxacin-resistance ^[19,24,25]^. Furthermore, zoliflodacin exhibited low MICs (0.03 and 0.06mg/L) in two fully ceftriaxone and cefixime resistant isolates.

Zoliflodacin is a novel spiropyrimidinetrione bacterial DNA gyrase/ topoisomerase inhibitor, which prevents bacterial DNA biosynthesis and results in accumulation of double-strand cleavages through a mechanism distinct from that in fluoroquinolones ^[18,24,26]^. In our study, all the ciprofloxacin-resistant zoliflodacin-sensitive isolates displayed double mutations in GyrA and greater than 97% had an additional amino acid substitutions in ParC. In contrast to fluoroquinolones, zoliflodacin targets the GyrB subunit of type II topoisomerase DNA gyrase, and specific mutations in GyrB can result in increased resistance to zoliflodacin ^[24,25]^. However, we found that only 4/143 (2.8%) of gonococcal isolates with the highest MICs (0.125 and 0.25 mg/L) harbored a GyrB mutation and the resultant amino acid substitutions/insertions (S467N, V470I or 480A) in GyrB were not associated with frank resistance. This finding differs from selection (albeit at low frequency) of resistance mutations of zoliflodacin that occurs in GyrB *in vitro* with first- and second-step mutations (D429N, D429A or K450T) that result in zoliflodacin MICs between 0.25–8 mg/L^[24,25]^. A modest temporal shift in MICs to zoliflodacin took place. Mutations in *mtrR*, which result in overexpression of the MtrCDE efflux pump, can increase efflux of antimicrobials and reduce the susceptibility to many antimicrobials ^[26,27]^. The MtrCDE efflux pump can also influence susceptibility to zoliflodacin^[25]^. Few studies have examined the impact of *parE* mutations on quinolone resistance in *N. gonorrhoeae*^[28,29]^. Clinical gonococcal isolates with P439S amino acid substitutions in ParE did not result in a significant increase in MIC to ciprofloxacin^[30]^. To our knowledge, three of the four ParE mutations identified in our study have not been reported previously.

In conclusion, zoliflodacin demonstrated potent in vitro antibacterial activity against a recent collection of clinical gonococcal isolates from China(2014 to 2018), including isolates with high-level resistance to ciprofloxacin, azithromycin and extended spectrum cephalosporins. Zoliflodacin MICs shifted upward temporally in the five-year period in the absence of clinical use. These results confirm the lack of pre-existing clinical resistance to zoliflodacin. Continued monitoring of antimicrobial susceptibility of zoliflodacin, a promising new oral antibacterial agent, for the treatment of uncomplicated gonorrhea is warranted.

## MATERIALS AND METHODS

### Bacterial isolates

From January 2014 to December 2018, a total of 986 gonococcal isolates were collected from male patients with symptomatic urethritis (urethral discharge and/or dysuria) attending the STD clinic at the Institute of Dermatology, Chinese Academy of Medical Sciences, Nanjing, China. All men except one reported that they were heterosexual. Urethral exudates were collected with cotton swabs, then immediately inoculated onto Thayer-Martin medium (Zhuhai DL Biotech, China) and cultured in candle jars at 36°C for 24–48 h. Gonococcal isolates were identified by colonial morphology, Gram’s stain and oxidase testing and growth on GC chocolate agar base (Difco, Detroit, MI) supplemented with 1% IsovitaleX™ (Oxoid, USA). Gonococcal colonies were suspended in tryptone-based soy broth and frozen (−70°C) until used for antimicrobial testing.

### Antimicrobial susceptibility testing

Zolifodacin powder was provided by Entasis, Therapeutics, Waltham, MA. The minimum inhibitory concentrations (MICs; mg/L) of *N. gonorrhoeae* isolates to zoliflodacin, penicillin, tetracycline, ciprofloxacin, spectinomycin, azithromycin, cefixime and ceftriaxone were determined by the agar dilution method in accordance with the Clinical and Laboratory Standards Institute (CLSI) guidelines^[31]^. ATCC 49226, WHO reference strains F, G, L, O, and P were used as quality controls. The MIC ranges of zoliflodacin for quality control (QC) strain ATCC 49226 were 0.125-0.25mg/L in each antimicrobial susceptibility testing run in this study in accordance with the defined MIC QC ranges (0.06-0.5mg/L) for zoliflodacin^[32]^. Criteria for decreased susceptibility to ceftriaxone (MIC⩾0.125 mg/L) and cefixime (MIC⩾0.25 mg/L) were defined by WHO^[33]^. Using CLSI^[31]^ and EUCAST ^[34]^ (for azithromycin only) criteria, the following MIC breakpoints were used to ascertain resistance: ⩾128 mg/L, spectinomycin; ⩾2 mg/L, penicillin and tetracycline and ⩾1 mg/L, ciprofloxacin and azithromycin. The breakpoint for zoliflodacin of ⩾0.5 mg/L was utilized as previously described ^[20]^. Multi-drug resistant (MDR) *N. gonorrhoeae* was defined as decreased susceptibility or resistance to extended spectrum cephalosporins (ESCs), plus resistance to at least two of the following antimicrobials: penicillin; ciprofloxacin and azithromycin ^[35,36]^.

### Identification of gene mutations that resulted in amino acid substitutions in GyrA, GyrB, ParC and ParE

For gonococcal isolates with zoliflodacin MICs of 0.125mg/L and 0.25mg/L, mutations in the quinolone-resistance-determining regions (QRDR) of *gyrA, gyrB, parC* and *par*E genes were determined by PCR and DNA sequencing using primers described previously ^[37–39]^ (supplemental Table 1). Genomic DNA was extracted from gonococcal isolates using the Rapid Bacterial Genomic DNA Isolation Kit (DNA-EZ Reagents V All-DNA-Fast-Out, Sangon Biotech Co. Ltd, Shanghai). PCR amplification and sequencing of the genes were carried out by Nanjing Qingke Biotech Co. Ltd.

### Data Analysis

Chi-square (χ^2^) testing was used to compare the rate of resistance in different years and Chi-square test for linear trends was used to assess the change in the MICs and the proportion of isolates resistant to antibiotics. SPSS version 19.0 was used for statistical analysis; P<0.05 was considered statistically significant.

## ACKNOWLEDGEMENTS

We thank Dr. Unemo Magnus for providing WHO reference strains.This work was supported by the grants from the Chinese Academy of Medical Sciences Initiative for Innovative Medicine (2016-I 2M-3-021) and the U.S. National Institutes of Health (AI084048 and AI116969).

## REFERENCES

1. Unemo M, Shafer WM. 2014. Antimicrobial resistance in *Neisseria gonorrhoeae* in the 21st Century: Past, evolution, and future. Clin Microbiol Rev 27:587–613. https://doi:10.1128/CMR.00010-14.

2. World Health Organization (WHO). 2016. WHO guidelines for the treatment of Neisseria gonorrhoeae. WHO: Geneva, Switzerland, 2016; Available from: https://www.who.int/reproductivehealth/publications/rtis/gonorrhoea-treatment-guidelines/en/ Accessed January 6, 2020.

3. Workowski KA, and Bolan GA. Centers for Disease Control and Prevention. 2015. Sexually transmitted diseases treatment guidelines, 2015. MMWR Recomm Rep. 64 (RR-03):1–137.

4. Yin YP, Han Y, Dai XQ, Zheng HP, Chen SC, Zhu BY, Yong G, Zhong N, Hu LH, Cao WL, Zheng ZJ, Wang F, Zhi Q, Zhu XY, and Chen XS. 2018. Susceptibility of *Neisseria gonorrhoeae* to azithromycin and ceftriaxone in China: A retrospective study of national surveillance data from 2013 to 2016. PLoS Med. 15:e1002499. https://doi:10.1371/journal.pmed.13

5. Tanaka M, Furuya R, Kobayashi I, Kanesaka I, Ohno A, Katsuse AK. 2018. Antimicrobial resistance and molecular characterization of *Neisseria gonorrhoeae* isolates in Fukuoka, Japan, from 1996 to 2016. J Glob Antimicrob Resist. pii: S2213–7165(18)30227-3. https://doi:10.1016/j.jgar.2018.11.011

6. Lahra MM, Enriquez R, George CRR. 2020. Australian gonococcal surveillance programme annual report, 2018. Commun Dis Intell (2018). 44. https://doi:10.33321/cdi.2020.44.4.

7. Day MJ, Spiteri G, Jacobsson S, Woodford N, Amato-Gauci AJ, Cole MJ, Unemo M; Euro-GASP network. 2018. Stably high azithromycin resistance and decreasing ceftriaxone susceptibility in *Neisseria gonorrhoeae* in 25 European countries, 2016. BMC Infect Dis. 18:609. https://doi:10.1186/s12879-018-3528-4.

8. Sexually Transmitted Disease Surveillance, 2018, Gonorrhea. Centers for Disease Control and Prevention. https://www.cdc.gov/std/stats18/gonorrhea.htm

9. Whiley DM, Mhango L, Jennison AV, Nimmo G, Lahra MM. 2018. Direct Detection of *penA* Gene Associated with Ceftriaxone-Resistant *Neisseria gonorrhoeae* FC428 Strain by Using PCR. Emerg Infect Dis. 24:1573–1575. https://doi:10.3201/eid2408.180295.

10. Golparian D, Rose L, Lynam A, Mohamed A, Bercot B, Ohnishi M, Crowley B, Unemo M. 2018. Multidrug-resistant *Neisseria gonorrhoeae* isolate, belonging to the internationally spreading Japanese FC428 clone, with ceftriaxone resistance and intermediate resistance to azithromycin, Ireland, August 2018. Euro Surveill. 23. https://doi:10.2807/1560-7917.ES.2018.23.47.1800617.

11. Chen SC, Han Y, Yuan LF, Zhu XY, Yin YP. 2019. Identification of internationally disseminated ceftriaxone-resistant *Neisseria gonorrhoeae* strain FC428, China. Emerg Infect Dis. 25: 1427–1429. https://doi:10.3201/eid2507.190172.

12. Yéo A, Kouamé-Blavo B, Kouamé CE, Ouattara A, Yao AC, Gbedé BD, Bazan F, Faye-Ketté H, Dosso M, Wi T, Unemo M. 2019. Establishment of a gonococcal antimicrobial surveillance programme, in accordance with World Health Organization standards, in Côte d’Ivoire, Western Africa, 2014-2017. Sex Transm Dis. 46:179–184. https://doi:10.1097/OLQ.0000000000000943.

13. Fifer H, Natarajan U, Jones L, Alexander S, Hughes G, Golparian D, Unemo M. 2016. Failure of dual antimicrobial therapy in treatment of gonorrhea. N Engl J Med. 374:2504–2506. https://doi:10.1056/NEJMc1512757.

14. Eyre DW, Sanderson ND, Lord E, Regisford-Reimmer N, Chau K, Barker L, Morgan M, Newnham R, Golparian D, Unemo M, Crook DW, Peto TE, Hughes G 6, Cole MJ, Fifer H, Edwards A, Andersson MI. 2018. Gonorrhoea treatment failure caused by a *Neisseria gonorrhoeae* strain with combined ceftriaxone and high-level azithromycin resistance, England, February 2018. Euro Surveill. 23. https://doi:10.2807/1560-7917.

15. Whiley DM, Jennison A, Pearson J, Lahra MM. 2018. Genetic characterisation of *Neisseria gonorrhoeae* resistant to both ceftriaxone and azithromycin. Lancet Infect Dis. 18:717–718. https://doi:10.1016/S1473-3099(18)30340-2.

16. World Health Organization (WHO). Global priority list of antibiotic-resistant bacteria to guide research, discovery, and development of new antibiotics; WHO: Geneva, Switzerland, 2017; Available from: https://www.who.int/medicines/publications/global-priority-list-antibiotic-resistant-bacteria/en/ Accessed January 6, 2020

17. The Centers for Disease Control and Prevention. The national action plan for combating antibiotic-resistant bacteria. 2015. Available from :https://www.cdc.gov/drugresistance/pdf/national_action_plan_for_combating_antibotic-resistant_bacteria.pdf. Accessed Febuary 10,2020.

18. Huband MD, Bradford PA, Otterson LG, Basarab GS, Kutschke AC, Giacobbe RA, Patey SA, Alm RA, Johnstone MR, Potter ME, Miller PF, Mueller JP. 2015. In vitro antibacterial activity of AZD0914, a new spiropyrimidinetrione DNA gyrase/topoisomerase inhibitor with potent activity against Gram-positive, fastidious Gram-Negative, and atypical bacteria. Antimicrob Agents Chemother 59:467–474. https://doi:10.1128/AAC.04124-14.

19. Jacobsson S, Golparian D, Alm RA, Huband M, Mueller J, Jensen JS, Ohnishi M, Unemo M. 2014. High in vitro activity of the novel spiropyrimidinetrione AZD0914, a DNA gyrase inhibitor, against multidrug-resistant *Neisseria gonorrhoeae* isolates suggests a new effective option for oral treatment of gonorrhea. Antimicrob Agents Chemother 58:5585–5588.https://doi:10.1128/AAC.03090-14.

20. Taylor SN, Marrazzo J, Batteiger BE, Hook EW 3rd, Seña AC, Long J, Wierzbicki MR, Kwak H, Johnson SM, Lawrence K, Mueller J. 2018. Single-dose zoliflodacin (ETX0914) for treatment of urogenital gonorrhea. N Engl J Med. 379: 1835–1845. https://doi:10.1056/NEJMoa1706988.

21. Su XH, Wang BX, Le WJ, Liu YR, Wan C, Li S, Alm RA, Mueller JP, Rice PA. 2016. Multidrug resistant *Neisseria gonorrhoeae* from Nanjing are sensitive to killing by a novel DNA gyrase inhibitor, ETX0914 (AZD0914). Antimicrob Agents Chemother. 60:621–623.https://doi:10.1128/AAC.01211-15.

22. Sethi S, Sharma D, Mehta SD, Singh B, Smriti M, Kumar B, Sharma M. 2006. Emergence of ciprofloxacin resistant *Neisseria gonorrhoeae* in north India. Indian J Med Res. 123(5):707–10.

23. Unemo M, Ringlander J, Wiggins C, Fredlund H, Jacobsson S, Cole M. 2015. High in vitro susceptibility to the novel spiropyrimidinetrione ETX0914 (AZD0914) among 873 contemporary clinical *Neisseria gonorrhoeae* isolates from 21 European countries from 2012 to 2014. Antimicrob Agents Chemother. 59(9):5220–5. https://doi:10.1128/AAC.00786-15.

24. Alm RA, Lahiri SD, Kutschke A, Otterson LG, McLaughlin RE, Whiteaker JD, Lewis LA, Su X, Huband MD, Gardner H, Mueller JP. 2015. Characterization of the novel DNA gyrase inhibitor AZD0914: low resistance potential and lack of cross-resistance in *Neisseria gonorrhoeae*. Antimicrob Agents Chemother. 59(3):1478–86.https://doi:10.1128/AAC.04456-14.

25. Foerster S, Golparian D, Jacobsson S, Hathaway LJ, Low N, Shafer WM, Althaus CL, Unemo M. 2015. Genetic Resistance Determinants, In Vitro Time-Kill Curve Analysis and Pharmacodynamic Functions for the Novel Topoisomerase II Inhibitor ETX0914 (AZD0914) in *Neisseria gonorrhoeae*. Front Microbiol. 6:1377. https://doi:10.3389/fmicb.2015.01377.

26. Unemo M, Shafer WM. 2014. Antimicrobial resistance in *Neisseria gonorrhoeae* in the 21st century: past, evolution, and future. Clin Microbiol Rev. 27:587–613.https://doi:10.1128/CMR.00010-14.

27. Golparian D, Shafer W M, Ohnishi, M, and Unemo M. 2014. Importance of multi-drug efflux pumps in the antimicrobial resistance property of clinical multi-drug resistant isolates of *Neisseria gonorrhoeae*. Antimicrob. Agents Chemother. 58, 3556–3559. https://doi:10.1128/AAC.00038-14

28. Kern G, Palmer T, Ehmann DE, Shapiro AB, Andrews B, Basarab GS, Doig P, Fan J, Gao N, Mills SD, Mueller J, Sriram S, Thresher J, Walkup GK. 2015. Inhibition of *Neisseria gonorrhoeae* type II topoisomerases by the novel spiropyrimidinetrione AZD0914. J Biol Chem 290:20984–20994. http://dx.doi.org/10.1074/jbc.M115.663534.

29. Lindbäck E, Rahman M, Jalal S, Wretlind B. Mutations in *gyrA, gyrB, parC*, and *parE* in quinolone-resistant strains of *Neisseria gonorrhoeae*. APMIS. 2002; 110(9):651–7.

30. Soge OO, Salipante SJ, No D, Duffy E, Roberts MC. 2016. In vitro activity of delafloxacin against clinical *Neisseria gonorrhoeae* isolates and selection of gonococcal delafloxacin resistance. Antimicrob Agents Chemother. 60(5):3106–11. https://doi:10.1128/AAC.02798-15.

31. Clinical and Laboratory Standards Institute. 2018. Performance Standards for Antimicrobial Susceptibility Testing: Twenty-Eighth Informational Supplement M100. CLSI, Wayne, PA, USA. 18

32. Miller AA, Traczewski MM, Huband MD, Bradford PA, Mueller JP. 2019. Determination of MIC Quality Control Ranges for the Novel Gyrase Inhibitor, Zoliflodacin. J Clin Microbiol. 57(9):1–7. https://doi:10.1128/JCM.00567-19.

33. WHO. 2012. Global Action Plan to Control the Spread and Impact of Antimicrobial Resistance in Neisseria gonorrhoeae. Available from: https://www.who.int/antimicrobial-resistance/global-action-plan/en/ Accessed January 6, 2020

34. The European Committee on Antimicrobial Susceptibility Testing. 2015. Breakpoint tables for interpretation of MICs and zone diameters. Version 5.0, 2015. Available from: https://www.eucast.org/ast_of_bacteria/previous_versions_of_documents/ Accessed January 6, 2020

35. Tapsall JW 1, Ndowa F, Lewis DA, Unemo M. 2009. Meeting the public health challenge of multidrug- and extensively drug-resistant *Neisseria gonorrhoeae*. Expert Rev Anti Infect Ther. 7(7):821–34. https://doi:10.1586/eri.09.63.

36. Clifton S, Bolt H, Mohammed H, Town K, Furegato M, Cole M, Campbell O, Fifer H, Hughes G. 2018. Prevalence of and factors associated with MDR *Neisseria gonorrhoeae* in England and Wales between 2004 and 2015: analysis of annual cross-sectional surveillance surveys. J Antimicrob Chemother. 73(4):923–932. https://doi:10.1093/jac/dkx520.

37. Tanaka M, Nakayama H, Haraoka M, Saika T. 2000. Antimicrobial Resistance of *Neisseria gonorrhoeae* and High Prevalence of Ciprofloxacin-Resistant Isolates in Japan, 1993 to 1998. J Clin Microbiol 38(2): 521–525.

38. Deguchi T, Yasuda M, Nakano M, Ozeki S, Kanematsu E, Kawada Y, Ezaki T, Saito I. 1996. Uncommon Occurrence of Mutations in the gyrB Gene Associated with Quinolone Resistance in Clinical Isolates of *Neisseria gonorrhoeae*. Antimicrob Agents Chemother 40(10): 2437–2438.

39. Unemo M, Fasth O, Fredlund H, Limnios A, Tapsall J. 2009. Phenotypic and genetic characterization of the 2008 WHO *Neisseria gonorrhoeae* reference strain panel intended for global quality assurance and quality control of gonococcal antimicrobial resistance surveillance for public health purposes. J Antimicrob Chemother 63(6): 1142–1151.

